# Dynamic response in the larval geoduck clam proteome to elevated *p*CO_2_

**DOI:** 10.1101/613018

**Authors:** Emma Timmins-Schiffman, José M. Guzmán, Rhonda Elliott, Brent Vadopalas, Steven B. Roberts

## Abstract

Pacific geoduck clams (*Panopea generosa*) are found along the Northeast Pacific coast where they are significant components of coastal and estuarine ecosystems and the basis of a growing and highly profitable aquaculture industry. The Pacific coastline, however, is also the sight of rapidly changing ocean habitat, including significant reductions in pH. The impacts of ocean acidification on invertebrate bivalve larvae have been widely documented and it is well established that many species experience growth and developmental deficiencies when exposed to low pH. As a native of environments that have historically lower pH than the open ocean, it is possible that geoduck larvae are less impacted by these effects than other species. Over two weeks in larval development (days 6-19 post-fertilization) geoduck larvae were reared at pH 7.5 or 7.1 in a commercial shellfish hatchery. Larvae were sampled at six time points throughout the period for a in-depth proteomics analysis of developmental molecular physiology. Larvae reared at low pH were smaller than those reared at ambient pH, especially in the prodissoconch II phase of development. Competency for settlement was also delayed in larvae from the low pH conditions. A comparison of proteomic profiles over the course of development reveal that these differing phenotypic outcomes are likely due to environmental disruptions to the timing of molecular physiological events as suites of proteins showed differing profiles of abundance between the two pH environments. Ocean acidification likely caused an energetic stress on the larvae at pH 7.1, causing a shift in physiological prioritization with resulting loss of fitness.

## Introduction

The Pacific geoduck (*Panopea generosa*) is a burrowing hiatellid clam found in low intertidal and subtidal sediments throughout the Northeast Pacific coast, including the US (Alaska, Washington, California), Canada (British Columbia), and Mexico (North Baja Pacific Coast) (Coan et al. 2000, Vadopalas et al. 2010). This bivalve mollusk is the largest clam in the world, reaching an average weight of 0.68 kg at maturity, with specimens over 3 kg having been recorded (Goodwin and Pease 1987). Geoducks are also among the longest-lived organisms in the animal kingdom, with an average reproductive lifespan of 30 years (Sloan and Robinson 1984). These clams are also known to play key roles in maintaining ecosystem health by filter feeding and biodeposition, as well as being a food source for marine organisms, including sea otters, fishes, crabs, and sea stars (Newell, 2004; Straus et al. 2009). Besides its ecological importance, geoducks are the most economically important clam fishery in North America (Hoffman et al., 2000), and the target of a growing aquaculture industry in Puget Sound (Washington, US) and British Columbia, with estimated annual production valued at US$21.4 million (FAO 2012). Pacific geoduck clams are considered to be particularly vulnerable to ocean acidification due to their calcifying nature and biogeographical distribution. As marine calcifiers, geoduck rely on calcite and aragonite for shell secretion (Orr et al. 2005; Weiss et al. 2002), both of which become less biologically available as pH declines (Feely et al. 2008). Additionally, Puget Sound and marine waters just off the Washington coast are naturally at lower pH than those of other regions (Busch et al., 2013), and ocean acidification has already caused a local decrease in pH of 0.05–0.15 units since the Industrial Revolution (Feely et al., 2010). Despite the ecological and economic importance of this giant clam in North America, the potential effect of ocean acidification on the physiology of the Pacific geoduck at its different life stages remain elusive.

The physiological impacts of ocean acidification on marine shelled mollusks during early life stages have been addressed in a number of species (e.g, Timmins-Schiffman et al., 2013; Stumpp et al., 2013; Waldbusser et al., 2013; Frieder et al., 2016; Kapsenberg et al., 2018). It is generally observed that low pH affects somatic growth and shell production in juvenile shellfish and alters respiration, feeding and excretion rates, energy availability and overall fitness. For example, short-term exposure to acidified waters in laboratory and in field settings caused significant shell dissolution and perforation of Sydney rock oyster juveniles (Dove and Sammut 2007a,b). Calcification rates declined linearly with increasing levels of CO2 in juvenile blue mussels and Pacific oysters (Gazeau et al., 2007), and a similar effect was described in eastern oyster larvae (Miller et al., 2009). Acidic conditions also reduced clearance, ingestion and respiration rates in juvenile clams, suggesting a slower growth in a future high CO2 coastal ocean (Fernandez-Reiriz et al., 2011). In juvenile eastern oysters, elevated CO2 levels reduced tissue energy stores (glycogen and lipid) and had a negative impact on soft tissue growth, indicating energy deficiency (Dickinson et al., 2012). High CO2 levels in water increased juvenile mortality rates and inhibited both shell and soft-body growth in juvenile eastern oyster (Beniash et al., 2010). The authors also found that elevated CO2 concentrations resulted in higher standard metabolic rates after a 20-week exposure, likely due to the higher energy cost of homeostasis. Interestingly, the rate of oxygen consumption was reduced to 65% after 20 h of exposure to hypercapnia conditions in juvenile marine mussels, clearly indicating a drastic reduction in metabolic rate (Michaelidis et al., 2005), suggesting that metabolic response to hypercapnia in mollusks may be species-specific and/or depend on exposure duration. While these and other studies provide critical information on how changes in seawater chemistry driven by ocean acidification are altering the physiological performance and ecology of marine shelled mollusks, the molecular mechanisms and pathways underlying these effects are unknown.

Mass-spectrometry (MS)-based proteomics technology provides a large-scale, unbiased approach for examining relative abundances of all proteins present in given sample. The proteome is closer to the realized phenotype than the genome or the transcriptome. In a previous study, the response of Pacific oyster larval proteome to ocean acidification was surveyed using low-resolution, 2-dimensional electrophoresis coupled with mass spectrophotometry (Dineshram et al., 2012). The authors found that 18.7% of the proteins detected in control samples (379 proteins total) were inhibited after exposure to a lower pH, including a severe inhibition of proteins related to calcification and cytoskeleton production. Similarly, proteomic alterations were observed in Sydney rock oyster larvae exposed to low pH (Parker et al. 2011). A proteomics study of *Crassostrea hongkongensis* pediveligers exposed to ocean acidification found an increased abundance of proteins involved in the antioxidant response and translation, with decreases in levels of proteins involved in structural molecules (Dineshram et al. 2015). Using a different proteomic approach, Spencer et al. (2019) evaluated the effect of natural pH variation on the proteome in juvenile geoduck. In a targeted proteomics assay of 13 environmentally sensitive proteins, no difference in abundance was found between habitats/pH regimes, suggesting that juvenile geoduck may tolerate a wide range of pH. However, like other invertebrate larval stages, geoduck larvae are likely highly sensitive to the effects of ocean acidification, which could have wide-ranging ecological and economic impacts. The effect of ocean acidification during earlier, more sensitive developmental stages or at other physiological processes (e.g., metabolism, biomineralization, cytoskeleton) remain unknown.

In this study, we use a high-resolution mass spectrometry-based proteomics data dependent analysis to (1) characterize for the first time the proteome of geoduck during early life stages (6 through 17 days post-fertilization) and (2) evaluate the physiological impact of a lower environmental pH at the proteome level. In depth coverage of the geoduck proteomes allowed us to establish that molecular hallmarks of developmental stage are not maintained across all physiological processes.

## Methods

### Larval Rearing

Broodstock maturation, spawning, and early larval development (days 1-4 post-fertilization) occurred in seawater treated with Na_2_CO_3_ (pH 8.2) in a commercial hatchery in Hood Canal, WA per normal operation conditions. At day 5 post-fertilization, larvae were exposed to ambient pH (7.5) or decreased pH conditions at 7.1 (2 conical tanks (200L) for each conditions). The pH of 7.5 is the ambient pH of incoming water to the hatchery and was achieved by removing the Na_2_CO_3_ treatment. A pH of 7.1 was obtained by removing Na_2_CO_3_ treatment and adding CO_2_ with a solenoid valve and venturi injector with a 5200 YSI probe for monitoring. Water was maintained at approximately 14°C throughout the experiment which ended 19 days post-fertilization (dpf). Initial stocking density of conical tanks (200L) was 14 million fertilized eggs.

Larvae were sampled on days on days 6, 8, 10, 12, 14, and 17 post-fertilization for larval proteomic analysis. Specifically, approximately ~10,000 larvae from each tank (two at pH 7.1 and two at pH 7.5) were rinsed with with isopropyl alcohol and stored at −80°C. For each sampling event larvae were measured in size classes using mesh screens. Number of larvae was estimated by weight based on an established conversion developed in the hatchery (Supporting Information 1). Larval settlement competency was determined when at least 30% of foot extension and movement was present in at least 30% of larvae on the screen.

The effect of pH on larval size was assessed used a Chi square test for homogeneity with a p-value cut-off of 0.05. Larval counts (calculated from larval mass) were averaged across conical tanks. Differences in larval count data among days was assessed via Chi square for days 10 and 17. Larvae were visually inspected for abnormalities on day 19.

### Protein extraction and LC-MS/MS

Larval proteins were digested and peptides were desalted following Timmins-Schiffman et al. (2014). Larval peptides were analyzed on an Orbitrap Fusion Lumos mass spectrometer (Thermo Scientific, Waltham, MA, USA) with a 4.5 cm, 100 μm pre-column and a 26 cm, 75 μm analytical column, both packed in-house with 3 μm C18 Dr. Maisch (Germany) beads. The analytical column was housed in a 50°C column heater for the duration of the analysis to improve chromatography. Over a 120 minute method, a 90-minute acetonitrile gradient went from 5-30%. In MS1 analysis in the Orbitrap, the resolution was 120K, scan range was 375-1575 *m/z*, max injection time was 50 ms, and AGC target was 700000. During the MS2 analysis in the IonTrap maximum injection time was 100 ms, AGC target was 2000, and centroid data was collected.

### Proteomics analysis

All mass spectrometry raw files were searched against a deduced geoduck larvae proteome with the addition of common laboratory contaminants from bovine, human, and other (cRAPome) (Supporting Information 2). Details on reference proteome development are described elsewhere (Supporting Information 3, https://github.com/sr320/paper-Pg-larvae-2019). Comet 2016.01 rev. 3 (Eng et al., 2012 and 2015) parameters included concatenated decoy search, mass tolerance of 20 ppm, 2 allowed missed trypsin cleavages, fragment bin tolerance of 1.0005 and offset of 0.4. Peptide and Protein Prophet (Keller et al., 2002; Nesvizhskii et al., 2003) were run consecutively using xinteract with no probability cut-off to allow for FDR cut-off later in the pipeline. Resulting pep.xml files were analyzed in Abacus (Fermin et al., 2011) with a FDR cut-off of 0.01 (probability from combined prot.xml file of 0.9) (Supporting Information 4).

Nonmetric multidimensional scaling (NMDS) analysis was performed on all proteins that were inferred across technical mass spectrometry replicates with at least 2 unique peptide spectral matches. Protein abundance data (normalized spectral abundance factor) were log(x+1) transformed and a Bray-Curtis dissimilarity matrix was calculated in R (R Core Team 2016). Eigenvectors were generated using envfit in the biostats package (McGarigal 2009). An eigenvector was considered significant if its axis loading value was >0.9 and its p-value was <0.001. Proteins corresponding to significant loadings on each axis in each NMDS were subjected to enrichment analysis following Timmins-Schiffman et al. (2017) at the following project-specific portal: https://meta.yeastrc.org/compgo_emma_geoduck_larvae/pages/goAnalysisForm.jsp. ANOSIM was performed on NSAF data standardized by row to assess significance of trends observed on the NMDS based on day and pH.

Differentially abundant proteins between consecutive time points (e.g., day 6 vs day 8 within pH condition) were determined using QSpec (Choi et al., 2008). Due to potential differences in developmental rate, differential protein abundances between treatments were not analyzed. For each QSpec analysis, only proteins that had a non-zero sum of spectral counts across replicates were included and spectral counts were summed across technical replicates.

In order to identify protein groups that had roles at specific time points during development, proteins with similar abundance trends were grouped using a hierarchical clustering method within each pH treatment. NSAF values were averaged across conical replicates within days and a Bray-Curtis dissimilarity matrix was calculated in R to use with the average clustering method. Cluster division was based on dendrogram topology (cut-off of h=0.5). Clusters were further classified into five categories based on the general trends through time determined with lowess smoothing of each cluster. Clusters including at least 10 proteins were subjected to enrichment analysis following Timmins-Schiffman et al. (2017). All R code used for analyses is provided at https://github.com/sr320/paper-Pg-larvae-2019 (Supporting Information 5).

## Results

### Larval Phenotype

At day 10 of the experiment, there was no observed difference in larval abundance between treatments, both having 2.3 million larvae (Figure 1). There was a difference in proportion of larvae in different size classes at day 10 (p<2.2e-16), due to more larvae in the <90 μm size class at pH 7.5 than at pH 7.1, but overall larval size was comparable at this time point between treatments (Supporting Information 6). By day 17 there was a clear difference in phenotype with larvae at pH 7.5 attaining a larger size by occupying larger size classes (180 μm and 200 μm) than larvae at pH 7.1, which did not exceed 180 μm (p<2.2e-16) (Figure 1). However, the larvae at pH 7.5 experienced a significant mortality by day 17 - 29 fold increased mortality compared to larvae at pH 7.1 (data now shown). Ciliates were detected at pH 7.5 at days 10 and 14 post-fertilization, but not at pH 7.1. At day 19, settlement was observed for larvae at pH 7.5 but not for larvae at pH 7.1.

**Figure 1.**
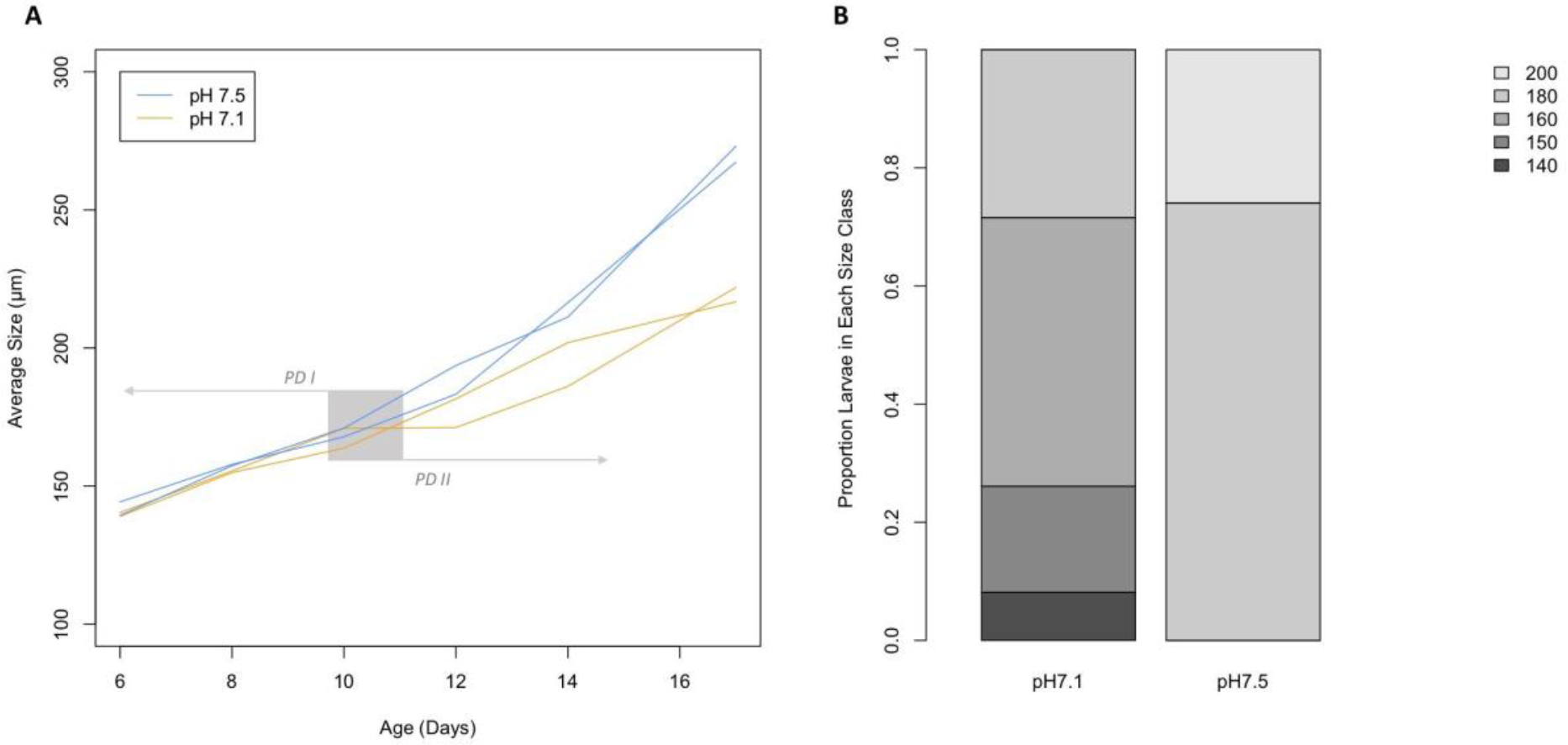
Growth curves of larvae from day 6 through 17 of development (A) and proportion of larvae in each size class (μm) on day 17 at pH 7.1 and 7.5 (B). Each individual tank in which larvae were reared is represented by a single line in panel A. Larval growth curves in tanks at pH 7.1 are orange and pH 7.5 tanks are in blue. The gray area indicates the larval transition from prodissoconch I (PD I) to II (PD II) (Goodwin & Pease 1989).

Across days 6 through 17 post-fertilization and at pH 7.5 and 7.1, 6328 proteins with at least 2 unique peptides across all biological and technical replicates were included in the dataset (Supporting Information 7). In examining global protein abundance there was a clear influence of time with significant separation of larval proteomes by day (ANOSIM R = 0.6579, p = 0.001), but not by pH (R = 0.008207, p = 0.338) (Figure 2). Proteomes are separated by time mostly along axis 1 of the NMDS, therefore, proteins with significant, positive eigenvector loadings along this axis (Supporting Information 8) play a dominant role in proteome differentiation over developmental time. The proteins with positive loadings are enriched for GO Biological Processes carbohydrate metabolic process, cellular amino acid metabolic process, and monocarboxylic acid metabolic process (https://github.com/sr320/paper-Pg-larvae-2019, Supporting Information 9).

**Figure 2.**
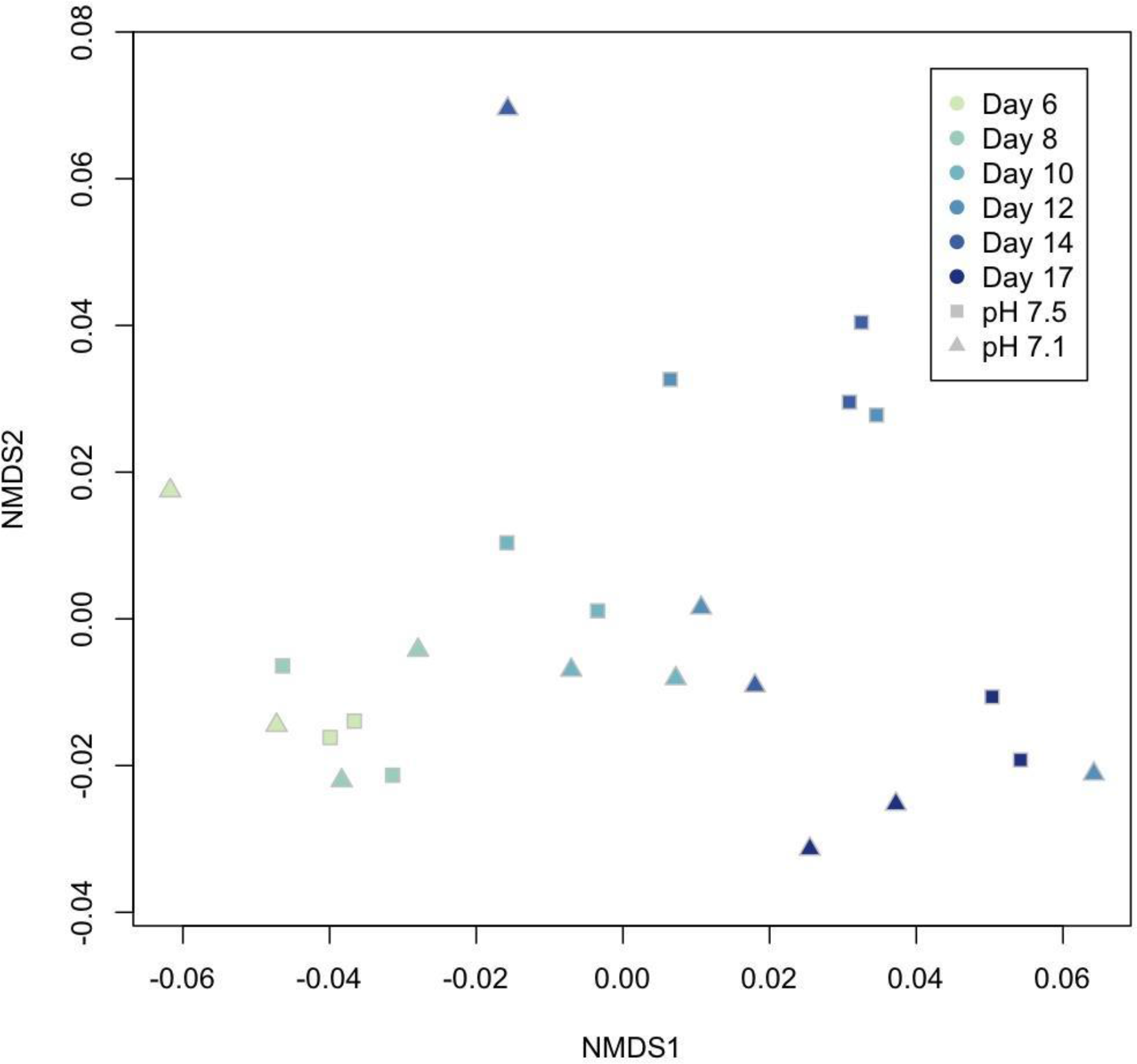
NMDS of larval proteomes across time and pH treatments. Increasingly dark colors correspond to progression through time (day 6 - 17). Squares represent proteomes from larvae reared at pH 7.5 and triangles represent larvae at pH 7.1.

The proteomic data from larvae reared at pH 7.5 clustered into 51 groups (43 of which had at least 10 protein members) and larvae reared at pH 7.1 clustered into 48 groups (40 of which had at least 10 protein members) based on abundance patterns over time (day 6 through 17) (Figures 3 and 4; Supporting Information 7). Cluster topology determined with lowess smoothing resulted in five types of abundance patterns over days 6-17 post-fertilization: stable/flat line (A), multiple peaks in abundance (B), single peak in abundance (C), general decrease over time (D), or general increase over time (E) (Table 1).

**Table 1.** Number of proteins that fall into each cluster type (A-E) across clusters at each pH. Proteins that are in clusters of fewer than 10 protein members are not categorized in a cluster type and so are not included.

| *Cluster Type* | *Number of Proteins pH 7.5* | *Number of Proteins pH 7.1* |
| --- | --- | --- |
| *A: stable/flat line* | 3678 | 4601 |
| *B: multiple peaks in abundance* | 1242 | 949 |
| *C: single peak in abundance* | 651 | 759 |
| *D: general decrease over time* | 1339 | 292 |
| *E: general increase over time* | 391 | 268 |

**Figure 3.**
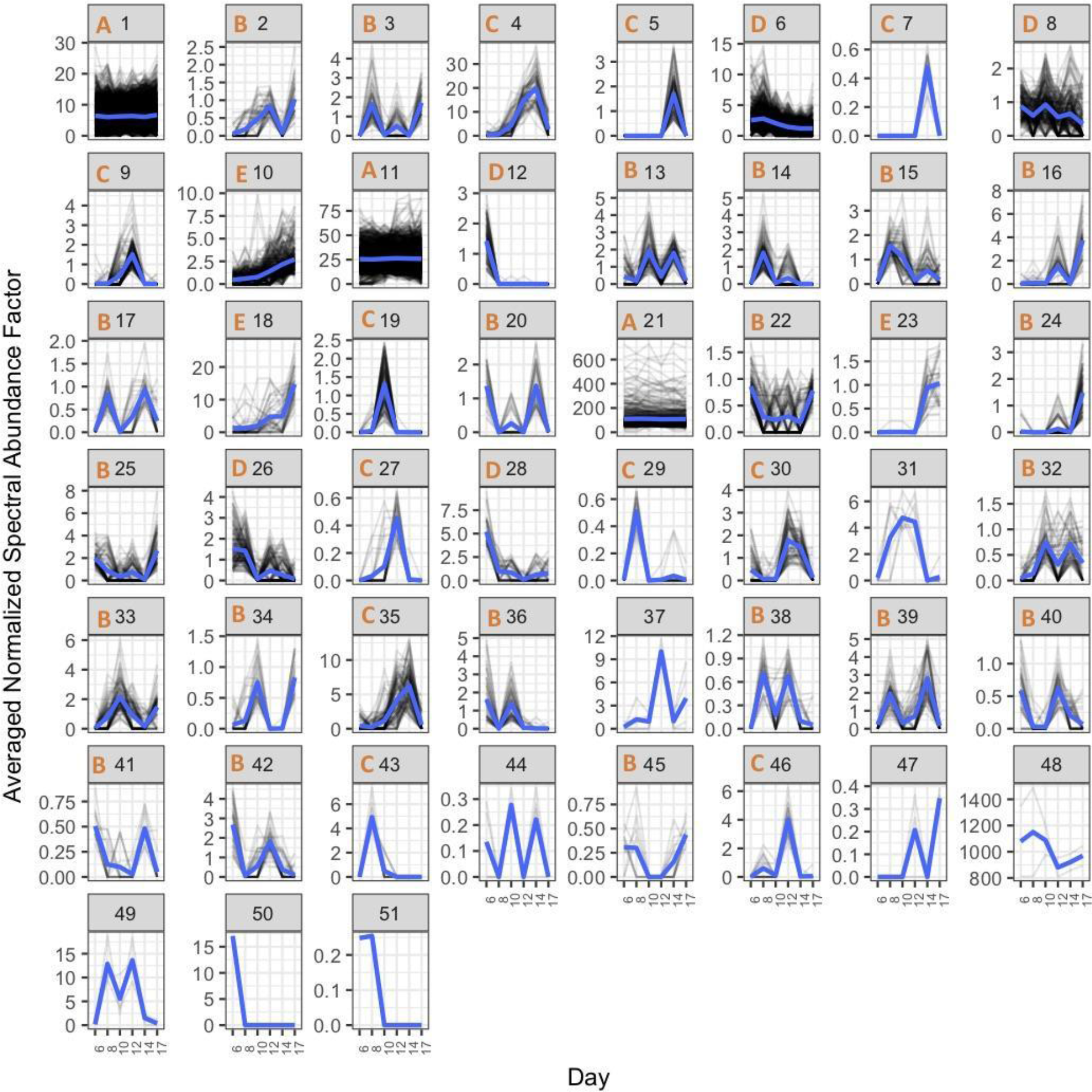
Plots of protein abundances of all proteins detected in pH 7.5-reared larvae from days 6-17. Proteins are divided among plots based on hierarchical clustering of abundance patterns across time, with normalized spectral abundance factor on the y-axis and time (days) on the x-axis. Lowess smoothed curves are represented by blue lines and abundance topology type is indicated by the orange letter at the top of each plot window for clusters with at least 10 protein members. Cluster number is indicated at the top of each plot window in black.

**Figure 4.**
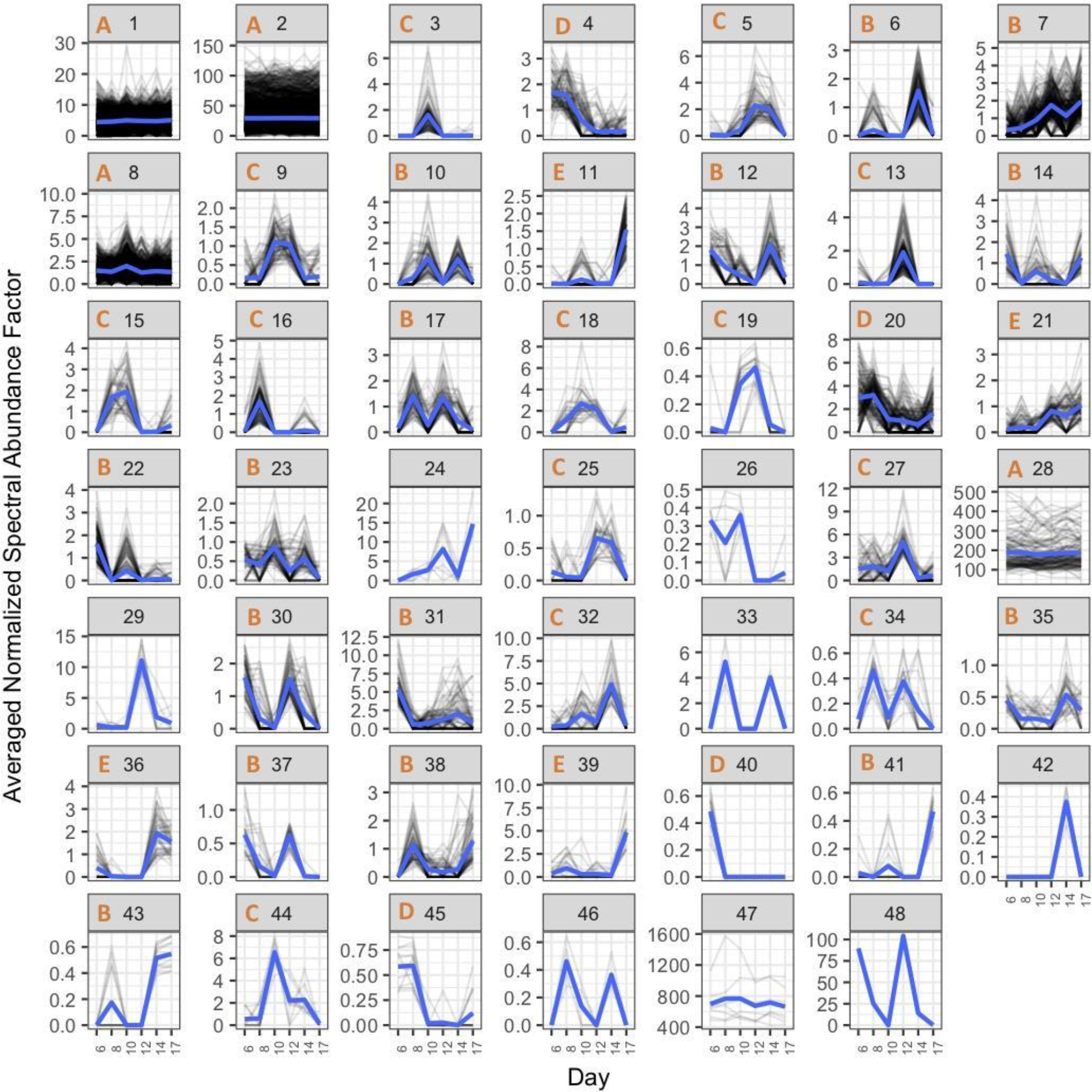
Plots of protein abundances of all proteins detected in pH 7.1-reared larvae from days 6-17. Proteins are divided among plots based on hierarchical clustering of abundance patterns across time, with normalized spectral abundance factor on the y-axis and time (days) on the x-axis. Lowess smoothed curves are represented by blue lines and abundance topology type is indicated by the orange letter at the top of each plot window for clusters with at least 10 protein members. Cluster number is indicated at the top of each plot window in black.

In an effort to define biological processes influenced by pH, proteins were grouped into three categories: (1) proteins that show stable abundance across time and are not altered by pH; (2) proteins that change in abundance during development but show similar trends at pH 7.5 and 7.1; and (3) proteins that change in abundance during development in patterns specific to a pH treatment.

### Basic Cellular Housekeeping is Maintained Across pH

Many of the protein functional groups that correspond to molecular housekeeping functions and that showed stable abundances across time (abundance pattern A) were maintained across both pH. The protein clusters that contain “housekeeping” proteins had high percentages (>50%) of shared protein members across pH treatments. These protein categories can be loosely described as ribosomes, translation, actin and myosin, proton-transporting ATP synthase, protein folding, and tricarboxylic acid cycle. For example, proteins from pH 7.5 cluster 21 were associated with the GO terms such as small ribosomal subunit (cellular component - CC), myosin complex (CC), proton-transporting ATP synthase complex catalytic core F(1) (CC), actin filament binding (molecular function - MF), unfolded protein binding (MF), translation (biological process - BP), tricarboxylic acid cycle (BP), and cell redox homeostasis (BP) representing 122 unique proteins. These proteins generally had high abundances and likely represent the core housekeeping proteins that allow other physiological processes to function.

At pH 7.1, cluster 1 proteins had mostly stable abundances and were enriched for the terms intracellular transport and cellular protein localization, which included nuclear transporters importin and exportin and cytoskeletal anchoring proteins. Cluster 2 proteins also appear to be relatively stable across time and the group was enriched for a diverse set of functions, such as cell redox homeostasis (e.g., thioredoxins, peroxiredoxins, glutathione reductase), cellular amino acid metabolic process (61 proteins involved in diverse amino acid metabolism processes), homophilic cell adhesion via plasma membrane adhesion molecules (e.g., protocadherins and cadherins), mucus layer (8 isoforms of mucin-5B), proteasome core complex alpha-subunit complex (e.g., proteasome complex subunit alpha, proteasome endopeptidase complex), and oxidoreductase activity.

### Some Stage Specific Proteins Are Robust to pH

Many protein groups demonstrated time-specific patterns of abundance, suggesting discrete roles at specific developmental time points. Our analysis allowed for comparison of broad protein functional categories across abundance patterns to discern trends of physiological activity at different time points. Many of these development-specific patterns were shared between pH 7.5 and 7.1 on the broad scale of functional category, suggesting essential molecular checkpoints that are resistant to alteration during a low pH response.

At day 10 post-fertilization, there was a signal of an up-regulation of cytoskeletal proteins at both pH 7.5 and 7.1. At pH 7.5, the proteins at higher abundance on day 10 were driven by the changing abundances of several tubulins. These tubulins drive the enrichment of the GO terms microtubule, GTP binding, GTPase activity, and structural constituent of cytoskeleton in the differentially abundant proteins (DAPs) that were elevated on day 10 compared to day 8. There were also two dynein proteins that were at higher abundance in this group. In pH 7.1-reared larvae, cytoskeletal proteins were at higher abundance on day 10 compared to both days 8 and 12, with the term structural constituent of cytoskeleton (including proteins such as tubulins and spectrin beta chain) in the comparison with day 8 and dynein complex (which included ciliary and axonemal dynein beta chains) in the comparison with day 12.

Day 12 post-fertilization emerges as an important physiological transition period based upon the molecular data and the phenotypic data since this is around the time of transition to prodissoconch II. Based on the proteomic data at both pH, this physiological transition is rooted in cell division, transcription, and translation. Proteins supporting DNA replication were at increased abundance at day 12. In cluster 46 (pH 7.5), the proteins contributing to the enriched term chromatin were histone H3 and a putative DNA helicase INO80 complex-like protein 1. Similarly, DAPs were involved in DNA replication and were at increased abundance at pH 7.5 on day 12 compared to 14 (DNA replication licensing factor MCM7, DNA helicase, and pre-mRNA-processing factor 19-like). At pH 7.1, the DAPs were enriched for methyltransferase activity (in a comparison with abundance at day 10) with proteins such as betaine--homocysteine S-methyltransferase 2, histone-arginine methyltransferase CARM1 and terms related to telomere maintenance and compared to day 14 with proteins such as X-ray repair cross-complementing protein 5 and ATP-dependent DNA helicase II subunit 2.

Transcription and translation also seemed to be up-regulated around the day 12 time point. At pH 7.5, proteins in the enriched term nuclear part (cluster 25) were mostly involved in transcriptional regulation and peaked in abundance on day 12. Protein glycosylation proteins also peaked on day 12 at pH 7.5 (cluster 2, two isoforms of polypeptide N-acetylgalactosaminyltransferase and galactosylgalactosylxylosyl protein 3-beta-glucuronosyltransferase), along with increased abundance of many ribosomal and other translation-related proteins on day 12 vs day 10. At pH 7.1 in clusters 13, 21, and 27, proteins with peaks in abundance on day 12 contributed to the enrichment of terms translation, ribosome, translational elongation, and others. The proteins that contributed to these terms included ribosomal proteins, a eukaryotic translation initiation factor, and elongation factors. There is also evidence of an increase in transcriptional activity, with proteins peaking on day 12 (and previously on day 6) in cluster 30 enriched for the term transcription factor activity sequence specific DNA binding.

There was likely more cellular growth and an increase in shell deposition around day 14 post-fertilization since larvae at both pH saw an increase in abundance of proteins involved in these processes. At pH 7.5, GO terms enriched in the proteins that were elevated on day 14 compared to day 17 included chitin metabolic process and chitin binding, which encompassed many IgGFc-binding proteins, chondroitin proteoglycan 2, chitotriosidase, and a chitinase. Many microtubule proteins were also included in this group (dyneins and coiled-coil domain proteins), contributing to the enrichment of terms microtubule-based movement, dynein complex, and microtubule motor activity. Like pH 7.5, DAPs that were elevated on day 14 at pH 7.1 included proteins involved in calcification and the cytoskeleton. Compared to day 12, DAPs elevated at day 14 were enriched for the terms mucus layer (3 isoforms of mucin-5B) and cell-matrix adhesion (sushi, nidogen, and EGF-domain containing protein 1; two forms of ependymin-related protein). Compared to day 17, proteins at increased abundance on day 14 were again enriched for mucus layer (4 forms of mucin-5B), chitin metabolic process (e.g., igGFc-binding protein, chitotriosidase), and dynein complex (e.g., dynein beta chain, ciliary; dynein gamma chain, flagellar outer arm).

DNA replication proteins again played an important role at the last time point: day 17 post-fertilization. Many DNA replication proteins were at increased abundance at pH 7.5, compared to day 14, perhaps suggestive of an upregulation of cell growth at day 17. These proteins include DNA helicases, DNA replication licensing factor MCM7, and X-ray repair cross-complementing protein 5, all of which contributed to the enrichment of terms such as DNA replication initiation, DNA duplex unwinding, and others. At pH 7.1, the GO term centrosome was enriched in cluster 41 proteins, which included two proteins that were detected only on day 17: coiled-coil domain containing protein 66 and centrosomal protein of 295 kDa. The pH 7.1 DAPs (compared to day 14) were enriched for DNA replication terms such as DNA conformation change (contributed to by proteins heterochromatin protein 1 binding protein 3, X-ray repair cross-complementing proteins 5, and DNA topoisomerase). There was also evidence of an up-regulation in transcription with the the proteins in pH 7.1 cluster 14 enriched for the term regulation of transcription, DNA-templated (see day 6) and the DAPs enriched for the term nucleic acid binding, examples of which are serine/arginine-rich splicing factor 4 and apoptotic chromatin condensation inducer in the nucleus.

### Developmental Physiology Can Be Disrupted by pH

Many molecular physiological processes were disrupted between pH 7.5 and 7.1 either as a result of the decreased pH and/or as an result of the probable developmental delay between the two pH. At every time point, there were proteomic signals that were specific to a pH, suggesting environmental-specific physiological needs in the two different environments.

#### Proteomic trends specific to pH 7.5

At day 6, there was proteomic evidence of changes to larval cytoskeletal and membrane structures at pH 7.5. There was an increase in proteins associated with the GO term spindle pole (cluster 42 proteins calmodulin and katanin p60 ATPase-containing subunit A). Three cytoskeletal DAPs were at increased abundance compared to day 8 (spectrin, dystonin, and stabilizer of axonemal microtubules) along with an endophilin. The GO molecular function (MF) term phospholipid binding was enriched in the DAPs (n=27 proteins), represented by 3 spectrin beta chains.

On day 8, some of the predominant molecular functions that arose at pH 7.5 were post-translational modifications of proteins, cilia biogenesis, and transport across cell membranes. Three protein abundance clusters have peaks in abundance at day 8. In cluster 3, proteins were enriched for terms associated with post-translational modification (e.g., peptidyl-tyrosine dephosphorylation) and the proteins that contribute to these terms include phosphatases, a cytoplasmic tRNA 2-thiolation protein, and E3 ubiquitin-protein ligases, suggesting increased protein turnover. In cluster 39 two integrator complex proteins (involved in transcription of small nuclear RNA) contributed to the enriched terms snRNA processing and integrator complex. There were 26 proteins in cluster 14 that were associated with the GO term integral component of membrane that are involved in diverse signaling and transport processes. Among the DAPs, the GO term motile cilium was enriched, based upon the increased abundance (at day 8 compared to 10) of three intraflagellar transport proteins that are essential in cilium biogenesis. These trends all suggest continued cellular growth and trafficking of materials across cell membranes.

At day 10 post-fertilization there was a strong signal of a metabolic shift at pH 7.5. This shift was evidenced by an increase in abundance of proteins involved in glucose and fatty acid metabolism (compared to day 8), which includes alpha-1,4, glucan phosphorylase, glucanase, phosphoenolpyruvate carboxykinase, 2-hydroxyacyl CoA lyase, acyl-CoA dehydrogenase, and phosphoethanolamine N-methyltransferase. Protein degradation enzymes were also at elevated levels, with increased abundances of cathepsin, ubiquitin-conjugating enzyme, tripeptidyl-peptidase, leucine aminopeptidase. Proteins in clusters 19 and 32 had abundance peaks on day 10, with the former proteins enriched for the term phosphatidylinositol-3-phosphate binding and the latter enriched for hexokinase activity and glucose binding (driven by two phosphotransferases, which are too general to truly categorize in this context). In cluster 19, the enrichment was driven by sorting nexin-27, which recycles plasma membrane proteins, and WD repeat domain phosphoinositide-interacting protein 3, which delivers cytoplasmic material to the lysosome. At day 10, there may be an increased metabolism of exogenous materials and thus greater cell growth.

This apparent up-regulation of transcription and translation on day 12 may have supported other physiological processes at pH 7.5, such as carbohydrate metabolism, as evidenced by the enrichment of the term cellular glucan metabolic process in cluster 10 proteins, which increased in abundance over time. More nuanced cellular processes, such as membrane budding, may have also been supported by this up-regulation as seen in the proteins in cluster 9 that were part of the enriched term ESCRT I complex (C-myc promoter-binding protein, multivesicular body subunit 12B, and vacuolar protein sorting-associated protein 37B).

On day 14 at pH 7.5, there was evidence of a large up-regulation of transcription and ribosomal proteins. There were peaks in abundance of protein groups in clusters 4, 35, and 39 (cluster 39 also had abundance peaks on day 8). All of these clusters were enriched for GO terms related to translation. In cluster 4, 27 proteins were responsible for enriched terms such as translation and structural constituent of ribosome; all of these 27 proteins with annotations were ribosomal proteins. Similarly, in cluster 35, 27 proteins (all annotated proteins were ribosomal) contributed to the enriched terms translation, ribosomal subunit, and structural constituent of ribosome.

Among the DAPs that were at elevated levels on day 14, many were ribosomal and translation elongation factors. In a comparison with abundances on day 12, proteins that were elevated on day 14 were enriched for organonitrogen compound metabolic process and structural constituent of ribosome (a group that included mostly ribosomal proteins). Proteins that decreased in abundance from day 14 to 17 were enriched for translation, ribosome biogenesis, and other related terms, based upon the inclusion of ribosomal proteins and translation elongation factors. The transition between days 14 and 17 had by far the highest number of DAPs.

On day 17, there was a decrease in abundance of ribosomal proteins (clusters 4 and 35 and DAPs), but an increase in abundance of proteins involved in post-translational modification and in transport. Proteins in clusters 2 and 3 peaked at earlier time points and at day 17 and included enrichment for the GO terms Golgi membrane (protein glycosylation proteins) and terms associated with post-translational modifications, respectively. Cluster 24 proteins, which peak on day 17, were enriched for the term integral component of membrane, which included proteins active in the Golgi, a variety of transporters, proteins involved in neurotransmitter release, and others. Similarly, the terms membrane budding and 1-phosphatidylinositol binding were enriched in the proteins at increased abundance on day 17 and include phosphatidylinositol-binding clathrin assembly proteins and protein SEC13 homolog. Proteins in cluster 25 also had a peak on day 17 (and at earlier time points), suggesting an up-regulation of transcriptional regulation. Carbohydrate metabolism proteins in cluster 10 (endoglucanase, 1,4,-alpha-glucan-branching enzyme (2 isoforms), and glycogen debranching enzyme) continued the trend of increased abundance throughout the latter part of the experiment.

#### Proteomic trends specific to pH 7.1

The molecular physiological profile of larvae reared at pH 7.1 on day 6 diverges from larvae reared at pH 7.5, in which the proteomic profile was dominated by cytoskeletal and membrane stability proteins. At pH 7.1, there was a strong signal of transcription- and translation-related processes at this early time point, with many of the contributing proteins detected at high abundances at later points in development in pH 7.5 larvae. For example, the term large ribosomal subunit is enriched in cluster 4 proteins at pH 7.1 and includes three 60S ribosomal proteins, all of which are found in pH 7.5 clusters with marked peaks much later in development (clusters 35 and 4). Transcription-associated processes that were enriched in clusters with upward abundance trends on day 6 included regulation of transcription, DNA-templated (cluster 14; e.g., Myc protein, transcription elongation factor STP5) and transcription factor activity, sequence-specific DNA binding (cluster 30; e.g., transcriptional repressor NF-X1, histone acetyltransferase). Other clusters with peaks at this time point were enriched for functions associated with cellular signaling, such as enzyme linked receptor protein signaling pathway (cluster 22; e.g., proprotein convertase subtilisin/kexin type 5, epidermal growth factor receptor), neurotransmitter:sodium symporter activity (cluster 37; sodium- and chloride-dependent neutral and basic amino acid transporter B(0+), sodium- and chloride-dependent glycine transporter), and Notch signaling pathway (cluster 40; protein jagged-1a, neurogenic locus Notch protein). These trends suggest a slightly different pathway in early development for larvae reared at pH 7.1.

The proteins that increased in abundance on day 8 at pH 7.1 were mostly involved in DNA replication and translation. None of the proteins that came to the forefront for the pH 7.1 analysis had the same trend at pH 7.5. Cluster 16 proteins contributed to the enriched term negative regulation of nitrogen compound metabolic process and included TIMELESS-interacting protein (involved in DNA repair), DNA methyltransferase 1-associated protein 1, and ATP-dependent RNA helicase Uap56. Similarly, the term helicase activity was enriched in the DAPs that were higher at day 8 than 6, which included RNA helicases and DNA helicase.

There was an increase in abundance of protein transport proteins at pH 7.1 on day 10. Larvae at day 10 (compared to 8) had higher abundances of proteins such as coatomer subunit alpha, protein transport protein SEC23, clathrin heavy chain, which led to the enrichment of the GO terms vesicle coat and transport vesicle. The proteins associated with a possible up-regulation of metabolism at pH 7.5 were absent in the analysis at pH 7.1

On day 12 post-fertilization there was a trend towards increases in abundance of proteins involved in metabolic processes in larvae at pH 7.1. Several proteins associated with biosynthetic processes contribute to the enriched term single-organism biosynthetic process in cluster 5 (e.g., kynureninase, enhancer of rudimentary homolog, and guanylate cyclase). In cluster 7, proteins that increased in abundance on day 12 contributed to the enriched terms phospholipid metabolic process (proteins include inositol monophosphatase 3, phospholipase, and ethanolamine phosphate transferase) and integral component of membrane (e.g., Voltage-dependent T-type calcium channel subunit alpha, transmembrane 9 superfamily member, V-type proton ATPase subunit A).

On day 17, cluster 43 proteins also peaked and were enriched for the term sodium ion transmembrane transporter activity based on the inclusion of sodium- and chloride-dependent glycine transporter and a sodium/hydrogen exchanger, the latter of which is involved in pH regulation.

## Discussion

A global, high-resolution proteomics survey detected the molecular hallmarks of specific developmental transition points throughout the geoduck larval period. Across environmental rearing conditions, the abundances of basic molecular housekeeping proteins (e.g., proteins related to translation, ATP synthase, etc.) were maintained at steady levels. Geoduck reared at different pH had different timing for developmental transitions. At pH 7.5, more proteins showed abundance trends that changed over time (types B-E), potentially suggesting more active regulation of physiological processes throughout development (Table 1). There were a higher number of proteins that did not change in abundance over days 6-17 post-fertilization (type A) at pH 7.1 than at pH 7.5, which could indicate less energy being allocated to specific physiological changes over time.

Across both pH treatments, there were several protein categories that maintained stable abundances throughout the developmental time frame captured and that can be considered essential, housekeeping proteins. These categories included ribosomes, myosin and actin, proton-transporting ATP synthase, protein folding, translation, and the citric acid cycle. In the clustered protein plots, these stably abundant protein clusters shared a higher percentage of proteins across pH treatments than other groups with time-specific abundance patterns, indicating the prioritization of maintenance of expression of thousands of proteins regardless of environmental pH.

The most dominant signal of proteomic differentiation across the experiment was based on developmental time, not on environmental pH, underlining the significance of the physiological shifts that occur during larval development. When considering proteomes across pH treatments, there was a trend of changes in abundance of proteins associated with various metabolic processes over time. This pattern suggests the importance of shifting larval metabolic needs over development.

### The molecular progression of development at pH 7.5

During early geoduck larval development (6-8 days post-fertilization) many molecular resources are dedicated to maintaining cellular integrity and growth. The cluster and differentially abundant protein (DAP) analyses both revealed higher abundances of cytoskeletal and membrane proteins at day 6 post-fertilization, which likely support cellular growth. By day 8, there was an increase in abundance of proteins involved in cilia biogenesis, transport across cell membranes, and post-translational modifications. The cilia biogenesis proteins could be a signal of general cellular growth and/or velum development. In other invertebrates, D-veliger larval proteomes had relatively higher abundances of proteins involved in cell division, shape, and cellular differentiation than trocophores, suggestive of development of more complex morphology and velum development (Huan et al., 2012; Di et al., 2017). Similarly, the abundance of cytoskeleton construction proteins increased from middle veliger through juvenile stages in the gastropod *Babylonia areolata* (Shen et al., 2018). These proteomic trends may indicate continued cellular growth during this high-growth early larval period, as well as protein turnover and cellular signaling and trafficking.

In the middle stages of larval development, there was continued evidence of cellular growth and suggestion of an up-regulation of metabolic processes. The cytoskeletal protein signal at day 10 was dominated by tubulins, suggesting that specific cytoskeletal proteins dominate at discrete developmental time points. Day 10 is also when there was a pronounced signal of glucose and fatty acid metabolism and protein degradation as the larvae break down algal resources for continued cellular growth and development. Proteins in cluster 10 (enriched for glucan metabolism) begin a steady increase in abundance through day 17 at day 10, suggesting continued carbohydrate metabolism to fuel growth and metamorphosis, a trend also observed in *B. areolata* (Shen et al., 2018). Ingestion rate increases throughout larval development in the geoduck *Panopea globosa* (Ferreira-Arrieta et al., 2015), which would be supported by upregulation of digestive and metabolic enzymes. Changes in abundances of digestion enzymes were also observed in *B. areolata* as feeding strategies shifted throughout development (Shen et al., 2018). By day 12 in geoduck, the increase in metabolic processes seems to support an up-regulation of transcription and translation, along with diverse other processes (e.g., transport, protein glycosylation, membrane budding).

Later developmental time points (days 14 and 17) in the geoduck mark a move towards the transition to the pediveliger stage and into competency for settlement and are characterized by molecular signals of translation, calcification, and protein turnover. On day 14 post-fertilization there was evidence of a large up-regulation of ribosomal proteins, suggesting increased protein translation at this time point. A similar signal was detected in the proteomes of competent larvae of the Pacific oyster, *Crassostrea gigas* (Huan et al., 2015). There was also evidence of an up-regulation of calcification in geoduck with increased abundances of several proteins associated with shell deposition. In the apple snail proteome, an increase in calcification-related proteins concurrent with greater rates of calcification was noted in the transition to becoming juvenile snails (Sun et al., 2010). By day 17, the ribosomal protein abundances had decreased, but were replaced with an increase in proteins involved in post-translational modifications and transport. There were also higher levels of proteins involved in neurotransmitter release and DNA replication on day 17, possibly indicative of greater cellular signaling and cell replication as the larvae underwent the major physiological shift into settled geoduck. Changes in expression of neurotransmitter pathway proteins and transcripts have been noted in other invertebrate larvae as they approach settlement (Niu et al., 2016; Shen et al., 2018).

### Impacts of low pH on geoduck larval development

Larvae reared at low pH (7.1) were significantly smaller than geoduck larvae reared at pH 7.5 at day 17 post-fertilization. This size discrepancy was observed earlier than day 17 and grew larger throughout development, with a relative flattening of the growth curve in low pH larvae compared to control counterparts. Invertebrate larvae reared at low pH have typically been observed to be smaller than those at higher pH (e.g., Kurihara et al., 2008; Parker et al. 2011; Andersen et al., 2013; Padilla-Gamino et al. 2013; Harney et al., 2016; Kapsenberg et al. 2018). Kurihara et al. (2008) targeted the beginning of pH-influenced developmental delay at the gastrula stage, which is when the shell field is first formed. In *C. gigas*, this difference in size has been observed to be larger in the prodissoconch II (PD II) phase (Timmins-Schiffman et al., 2013), as observed in the current study. In the mussel *Mytilus galloprovincialis*, larvae reared at low pH lacked the distinctive boundary between PD I and PD II shell material (Kurihara et al., 2008). It is possible that there the developmental transition from PD I to PD II (a transition from amorphous calcium carbonate to an aragonitic shell; Medavkovic et al., 1997; Seung Woo et al., 2006) at low pH is too energetically demanding to be executed successfully when the environment is not conducive to CaCO3 formation. Without information on developmental stage at the different time points, it is difficult to assess whether this slowed growth represents a delay in developmental rate or simply reduced size at low pH, both of which have been observed in invertebrate larvae. Given that larvae at pH 7.5 were larger and competent to settle before larvae reared at pH 7.1, the developmental delay hypothesis could be supported. In fact, metamorphosis in geoduck is known to be delayed by stress (Goodwin and Pease 1989). Developmental delay due to ocean acidification has been observed in other bivalve larvae, such as *C. gigas* and *M. galloprovincialis* (Kurihara et al., 2008; Timmins-Schiffman et al. 2013; Frieder et al., 2016; De Wit et al., 2018).

The other marked difference in larvae between the two pH treatments was a disconnect in timing of molecular development. This disconnect was also observed in *C. gigas* larvae reared at different pH and attributed to a potential delay in development (i.e. sampling of different developmental time points) (Harney et al., 2016). Whereas many housekeeping proteins maintained stable abundances across both pH treatments, the proteins that turned “on” and “off” at specific developmental time points differed (Table 1). When comparing proteins with shared membership between clusters at different pH, none of the non-housekeeping clusters shared >50% of their protein members, indicating discrete patterns of protein abundance based on environmental pH. For example, at pH 7.5 there was a large signal of ribosomal protein upregulation around days 12 and 14, but a similar signal occurred around days 6 and 8 at pH 7.1. These contrasts suggest distinct developmental paths based upon environmental conditions, with at least one phenotypic outcome being significantly smaller larval size.

Early developmental timepoints for geoduck reared at pH 7.1 were dominated by higher abundances of proteins involved in DNA replication, transcription, translation, and signaling. At day 6, there was a strong proteomic signal of ribosomal proteins, proteins that support transcription, and cellular signaling. Unhindered protein synthesis under low pH conditions has been observed in *C. gigas* (Frieder et al. 2016) and maintenance of translation-related proteins may be further evidence that ocean acidification does not impact this essential process. At day 8, the molecular developmental signals were still diverged between the two pH treatments, with an increase in abundance of DNA replication and translation proteins at pH 7.1.

Days 10 and 12 of development at pH 7.1 saw proteomic evidence of cytoskeletal modification, metabolism, and cellular growth. On day 10 at both pH, proteins involved in the cytoskeleton were at increased abundance suggesting important shifts in cellular morphology and/or growth at this time point. Increased pCO2 has a documented effect on the level of cytoskeletal gene transcripts, which decreased at elevated pCO2 in urchin pluteus larvae (Padilla-Gamino et al., 2013). Protein turnover also seems to be an important process at both pH, with protein transport and translation trending at pH 7.1 and protein degradation and post-translational modification proteins at increased abundances at pH 7.5. There were also signals of an up-regulation of metabolism at both pH, however this signal was more pronounced at days 10 and 12 at pH 7.5 and at day 12 only at pH 7.1, with increases in abundances of proteins involved in biosynthesis and phospholipid metabolism at the lower pH. Low environmental pH has been found to negatively affect sea urchin larvae digestion by decreasing the alkalinity of their stomachs (Stumpp et al. 2013), which could also impact geoduck and thus change the molecular signals of metabolism.

At days 14 and 17 post-fertilization there were shared signals across pH of proteins involved in calcification, cytoskeleton, and transcription, but some distinct molecular signals as well. There was strong signal of calcification-related proteins on day 14 post-fertilization at pH 7.1, with increased abundances of several mucin isoforms, IgGFc-binding protein, and chitotriosidase. These suggest an up-regulation of calcification in the prodissoconch II stage of development. By day 17, the increase in abundances of proteins involved in DNA replication, transcription, and translation may signal an up-regulation of cellular growth at pH 7.1 as the larvae approach the point of metamorphosis. A comparison of gene transcripts between low and high pH in sea urchin larvae revealed decreased abundance of genes involved in the nucleosome and chromatin organization and assembly (Padilla-Gamiño et al. 2013). Given that larvae from pH 7.5 were competent to settle two days after the last proteomic time point and low pH larvae were not, we can postulate that some of the proteomic signals detected at the higher pH (and not at pH 7.1) are necessary for reaching competency. Some of these protein categories not detected in our analysis of pH 7.1 were neurotransmitter release, membrane budding, and carbohydrate metabolism. It is possible that the low pH larvae were not in the appropriate metabolic state to send/receive the intercellular signals to begin metamorphosis.

## Conclusions

By following invertebrate larval development over two weeks, this data set reveals that larvae reared at low pH experience disruptions to their molecular physiology. These disrupted patterns observed in the proteome are likely a result of the energetic toll placed on the larvae as they grow and calcify at a low pH that may negatively impact multiple physiological processes (e.g., Waldbusser et al., 2013; Stumpp et al., 2013). Despite these significant impacts on many molecular pathways, essential, housekeeping proteins were maintained at stable levels, as observed at the higher pH. This maintenance of certain pathways over others provides insight into the possible prioritization of energetic resources in under stressful environmental conditions. However, these collective housekeeping pathways were not enough to see the larvae through to full competency and settlement as the low pH geoduck larvae were smaller, had higher levels of abnormal characteristics, were likely developmentally delayed, and were not competent to settle by day 19 post-fertilization. Thus, even though geoduck clams live in a naturally lower pH environment, their larval stages are not robust to the impacts of prolonged ocean acidification. As the largest biomass in Puget Sound, WA geoduck are integral parts of a local and more extensive ecosystem. As pH continues to drop, their populations may be at risk of lower larval survival and recruitment, with consequences for nutrient cycling and higher trophic levels.

## References

Ainsworth C.H., Samhouri J.F., Busch D.S., Chueng W.W.L., Dunne J., Okey T.A. (2011) Potential impacts of climate change on Northeast Pacific marine fisheries and food webs. ICES Journal of Marine Science 68: 1217–1229.

Andersen S., Grefsrud E.S., Harboe T. (2013). Effects of increased pCO2 level on early shell development in the great scallop (*Pecten maximus* Larmack) larvae. Biogeosciences, 10, 6161–6184.

Beardall J., Stojkovic S., Larsen S. (2009) Living in a high CO2 world: impacts of global climate change on marine phytoplankton. Plant Ecology & Diversity 2: 191–205

Beniash E., Ivanina A., Lieb N.S., Kurochkin I., Sokolova I.M. (2010). Elevated level of carbon dioxide affects metabolism and shell formation in oysters *Crassostrea virginica*. Mar. Ecol. Prog. Ser. 419, 95–108.

Busch D.S., Harvey C.J., McElhany P. (2013) Potential impacts of ocean acidification on the Puget Sound food web. ICES Journal of Marine Science 70: 823–833.

Cai W.J, Hu X., Huang W.J., Murrell M.C., Lehrter J.C., Lohrenz S.E., Chou,W.C., Zhai, W. Hollibaugh, J.T., Wang, Y., Zhao, P. Guo, X., Gundersen, K., Dai, M., Gong, G.C. (2011) Acidification of subsurface coastal waters enhanced by eutrophication. Nat Geosci. 2011; 4(11):766–70.

Caldeira K., and Wickett M.E. (2003) Anthropogenic carbon and ocean pH. Nature 425:365

Choi H., Fermin D., Nesvizhskii A.I. (2008). Significance analysis of spectral count data in label-free shotgun proteomics. Molecular and Cellular Proteomics, 7(12), 2373–2385.

Coan E.V., Scott P.V., Bernard F.R. (2000). Bivalve Seashells of Western North America: Marine Bivalve Mollusks from Arctic Alaska to Baja California. Santa Barbara Museum of Natural History

De Wit P., Durland E., Ventura A., Langdon C.J. (2018). Gene expression correlated with delay in shell formation in larval Pacific oysters (*Crassostrea gigas*) exposed to experimental ocean acidification provides insights into shell formation mechanisms. BMC Genomics, 19, 160.

Di G., Kong X., Miao X., Zhang Y., Huang M., Gu Y., You W., Zhang J., Ke C. (2017). Proteomic analysis of trocophore and veliger larvae development in the small abalone *Haliotis diversicolor*. BMC Genomics, 18, 809.

Dickinson GH, Ivanina AV, Matoo OB, Pörtner HO, Lannig G, Bock C, Beniash E, Sokolova IM (2012) Interactive effects of salinity and elevated CO2 levels on juvenile eastern oysters, *Crassostrea virginica*. J Exp Biol 215:29–43

Dineshram R, Wong KKW, Xiao S, Yu Z, Qian PY, Thiyagarajan, V. (2012) Analysis of Pacific oyster larval proteome and its response to high-CO2. Mar Pollut Bull 64: 2160–2167

Dineshram R., Quan Q., Sharma R., Chandramouli K., Yalamanchili H. K., Chu I., Thiyagarajan V. (2015). Comparative and quantitative proteomics reveal the adaptive strategies of oyster larvae to ocean acidification. Proteomics, 15(23-24), 4120–4134.

Dove M.C., Sammut J. (2007a) Impacts of estuarine acidification on survival, growth of Sydney rock oysters *Saccostrea glomerata* (Gould 1850). J Shellfish Res 26(2):519–527

Dove M.C., Sammut J. (2007b) Histological and feeding response of Sydney rock oysters, *Saccostrea glomerata*, to acid sulfate soil outflows. J Shellfish Res 26(2):509–518

Duarte C. M. Hendriks I.E., Moore T.M., Olsen Y.S., Steckbauer A., Ramajo L., Carstensen J., Trotter J.A., McCulloch M. (2013) Is ocean acidification an open-ocean syndrome? Understanding anthropogenic impacts on seawater pH. Estuar. Coasts 36, 221–236

Eng J.K., Jahan T.A., Hoopmann M.R. (2012). Comet: an open source tandem mass spectrometry sequence database search tool. Proteomics. doi: 10.1002/pmic.201200439.

Eng J.K., Hoopmann M.R., Jahan T.A., Egertson J.D., Noble W.S., MacCoss M.J. (2015). A Deeper Look into Comet - Implementation and Features. Journal of the American Society of Mass Spectrometry. doi: 10.1007/s13361-015-1179-x

Feely R.A., Sabine C.L., Hernandez-Ayon J.M., Ianson D., Hales B. (2008) Evidence for upwelling of corrosive ‘acidified’ water onto the continental shelf. Science 320: 1490–1492

Feely R.A., Alin S.R., Newton J., Sabine C.L., Warner M., Devol, A., Krems, C., Maloy, C. (2010) The combined effects of ocean acidification, mixing, and respiration on pH and carbonate saturation in an urbanized estuary. Estuarine, Coastal and Shelf Science 88: 442–449.

Fermin D., Basrur V., Yocum A.K., Nesvizhskii A.I. (2011). Abacus: a computational tool for extracting and pre-processing spectral count data for label-free quantitative proteomic analysis. Proteomics, 11(7), 1340–1345.

Fernández-Reiriz, M.J., Range, P., álvarez-Salgado, X.A., Labarta, U. (2011) Physiological energetics of juvenile clams *Ruditapes decussatus* in a high CO_2_ coastal ocean. Marine Ecology Progress Series 433: 97–105.

Ferreira-Arrieta A., García-Esquivel Z., González-Gómez M.A., Valenzuela-Espinoza E. (2015). Growth, survival, and feeding rates for the geoduck *Panopea globosa* during larval development. Journal of Shellfish Research, 34(1), 55–61.

Frieder C.A., Applebaum S.L., Pan T.C.F., Hedgecock D., Manahan D.T. (2016). Metabolic cost of calcification in bivalve larvae under experimental ocean acidification. ICES Journal of Marine Science, 74(4), 941–954.

Gazeau F., Quiblier C., Jansen J.M., Gattuso J.P., Middelburg J.J., Heip C.H.R. (2007). Impact of elevated CO2 on shellfish calcification. Geophys. Res. Lett. 34 (7).

Goodwin C.L., and Pease P. (1987). The distribution of geoduck (*Panope abrupta*) size, density, and quality in relation to habitat characteristics such as geographic area, water depth, sediment type, and associated flora and fauna in Puget Sound, Washington. State of Washington, Department of Fisheries, Shellfish Division.

Goodwin C.L., and Pease B. (1989). Species Profiles: Life Histories and Environmental Requirements of Coastal Fishes and Invertebrates (Pacific Northwest). Pacific Geoduck Clam. Biological Report 82 U.S. Army Corps of Engineers and U.S. Department of the Interior, 22 pp.

Gosselin, L. A., and Qian, P. (1997) Juvenile mortality in benthic marine invertebrates. Mar. Ecol. Prog. Ser. 146: 265–282.

Hall-Spencer J.M., Rodolfo-Metalpa R., Martin S., Ransome E. Fine M., Turne, S.M., Rowley S.J., Tedesco D., Buia M.C. (2008) Volcanic carbon dioxide vents show ecosystem effects of ocean acidification. Nature 454:96–99

Harney E., Artigaud S., Le Souchu P., Miner P., Corporeau C., Essid H., Pichereau V., Nunes F.L.D. (2016). Non-additive effects of ocean acidification in combination with warming on the larval proteome of the Pacific oyster, *Crassostrea gigas*. Journal of Proteomics, 135, 151–161.

Hauri C., Gruber N., Plattner G.K., Allin S., Feely R.A., Hales B., Wheeler P.A. (2009) Ocean acidification in the California Current System. Oceanography 22:60–71

Hoffmann A., Bradbury A., Goodwin C.L. (2000) Modeling geoduck, *Panopea abrupta* (Conrad, 1849) population dynamics. I. Growth. J. Shellfish Res. 2000, 19 (1), 57−62.

Huan P., Want H., Dong B., Liu B. (2012). Identification of differentially expressed proteins involved in early larval development of the Pacific oyster *Crassostrea gigas*. Journal of Proteomics, 75(13), 3855–3865.

Huan P., Wang H., Liu B. (2015). A Label-Free Proteomic Analysis on Competent Larvae and Juveniles of the Pacific Oyster *Crassostrea gigas*. PLoS ONE, 10(8), e0135008.

Kapsenberg L., Miglioli A., Bitter M.C., Tambutté E., Dumollard R., Gattuso J.P. (2018). Ocean pH fluctuations affect mussel larvae at key developmental transitions. Proceedings of the Royal Society B, 285(1893), 20182381.

Keller A., Nesvizhskii A.I., Kolker E., Aebersold R. (2002). Empirical statistical model to estimate the accuracy of peptide identifications made by MS/MS and database search. Analytical Chemistry, 74(20), 5383–5392.

Kelly R.P., Foley M.M., Fisher W.S., Feely R.A., Halpern B.S., Waldbusser G.G., Caldwell M.R. (2011). Mitigating local causes of ocean acidification with existing laws, Science, 332(6033), 1036–1037.

Kroeker K.J., Kordas R.L., Crim R.N., Singh G.G. (2010). Meta-analysis reveals negative yet variable effects of ocean acidification on marine organisms. Ecol. Lett. 13, 14191434.

Kurihara H., Takamasa A., Kato S., Ishimatsu A. (2008). Effects of elevated pCO2 on early development in the mussel *Mytilus galloprovincialis*. Aquatic Biology, 4, 225–233.

Medavkovic D., Popovic S., Grzeta B., Plaxonic M., Hrs-Brenko M. (1997). X-ray diffraction study of calcification processes in embryos and larvae of the brooding oyster *Ostrea edulis*. Marine Biology, 129(4), 615–623.

McGarigal K. (2009). BIOSTATS. R package. https://www.umass.edu/landeco/teaching/ecodata/labs/biostats.pdf

Michaelidis B., Ouzounis C., Paleras A., Pörtner, H. (2005). Effects of long-term moderate hypercapnia on acid–base balance and growth rate in marine mussels *Mytilus galloprovincialis*. Marine Ecology Progress Series. 293 119–118.

Miles H., Widdicombe S., Spicer J.I., Hall-Spencer J. (2007) Effects of anthropogenic seawater acidification on acid-base balance in the sea urchin *Psammechinus miliaris*. Mar Pollut Bull 54:89–96

Nesvizhskii A.I., Keller A., Kolker E., Aebersold R. (2003). A statistical model for identifying proteins by tandem mass spectrometry. Analytical Chemistry, 75(17), 4646–4658.

Newell R.I. (2004). Ecosystem influences of natural and cultivated populations of suspension-feeding bivalve mollusks: A review. J. Shellfish Res., 23 (1), 51−62.

Nilsson G.E., Dixson D.L., Domenici P., McCormick M.I., Sorensen C., Watson, S.A, Munday, P.L. (2012). Near-future carbon dioxide levels alter fish behaviour by interfering with neurotransmitter function. Nature Climate Change 2: 201–204.

Niu D., Wang F., Xie S., Sun F., Wang Z., Peng M., Li J. (2016). Developmental Transcriptome Analysis and Identification of Genes Involved in Larval Metamorphosis of the Razor Clam, *Sinonovacula constricta*. Marine Biotechnology, 28(2), 168–175.

Orr J.C., Fabry V.J., Aumont O., Bopp L., Doney S.C., Feel F.A., Gnanadesikan A., Gruber N., Ishida A., Joos F., Key R.M., Lindsay K., Maier-Reimer E., Matear R., Monfray P., Mouchet, A., Najjar R.G., Plattner G.K., Rodgers K.B., Sabine C.L., Sarmiento J.L., Schlitzer R., Slater R.D., Totterdell I.J., Weirig M.F., Yamanaka Y., Yool, A. (2005) Anthropogenic ocean acidification over the twenty-first century and its impact on calcifying organisms. Nature 437:681–686

Padilla-Gamiño J.L., Kelly M.W., Evans T.G., Hofmann G.E. (2013). Temperature and CO2 additively regulate physiology, morphology and genomic responses of larval sea urchins *Strongylocentrotus purpuratus*. Proceedings of the Royal Society B, 280(1759), 20130155.

Parker L., Ross P., Raftos D., Thompson E., O’Connor W. (2011). The proteomic response of larvae of the Sydney rock oyster, *Saccostrea glomerata* to elevated pCO2. Australian Zoologist, 35(4), 1011–1023.

Pechenik J.A. (1999). On the advantages and disadvantages of larval stages in benthic marine invertebrate life cycles. Mar Ecol Prog Ser 177: 269–297.

Pelejero C., Calvo E., Hoegh-Guldberg, O. (2010). Paleoperspectives on ocean acidification. Trends in Ecology & Evolution, 25: 332–344.

Pörtner H.O., Langenbuch M., Reipschläger A. (2004). Biological impact of elevated ocean CO_2_ concentrations: Lessons from animal physiology and earth history. Journal of Oceanography. 60(4), 705–718.

R Core Team (2016). R: A language and environment for statistical computing. R Foundation for Statistical Computing, Vienna, Austria. URL https://www.R-project.org/.

Raven J., Caldeira K., Elderfield H., Hoegh-Guldberg O., Liss P., Riebesell U., Shepherd J., Turley C., Watson A., Heap R., Banes R., Quinn R. (2005) Ocean acidification due to increasing atmospheric carbon dioxide. Royal Society Policy Doc 12/05, Clyvedon Press, Cardiff.

Salisbury J., Green M., Hunt, C., Campbell J. (2008). Coastal acidification by rivers: a threat to shellfish? Eos Trans Amer Geophys Union 89 (50) 513–513.

Shen M., Di G., Li M., Fu J., Dai Q., Miao X., Huang M., You W., Ke C. (2018). Proteomics Studies on the three Larval Stages of Development and Metamorphosos of *Babylonia areolata*. Scientific Reports, 8, 6269.

Sloan N.A., and Robinson S.M.C. (1984). Age and gonad development in the geoduck clam *Panope abrupta* (Conrad) form southern British Columbia, Canada. J. Shellfish. Res. 4: 131–137.

Spencer L.H., Horwith M., Lowe A.T., Venkataraman Y.R., Timmins-Schiffman E., Nunn B.L., Roberts S.B. (2019). Pacific geoduck (*Panopea generosa*) resilience to natural pH variation. Comparative biochem and Physiology Part D: Genomics and Proteomics. 30: 91–101.

Spicer J.I., Raffo A., Widdicombe S. (2007) Influence of CO2-related seawater acidification on extracellular acid–base balance in the velvet swimming crab Necora puber. Mar Biol 151:1117–1125

Straus K., Crosson L., Vadopalas B. (2009). Effects of Geoduck Aquaculture on the Environment: A Synthesis of Current Knowledge. School of Aquatic and Fishery Sciences, University of Washington. 64 pages.

Sun J., Zhang Y., Thiyagarajan V., Qian P.Y., Qui J.W. (2010) Protein expression during the embryonic development of a gastropod. Proteomics, 10(14), 2701–2711.

Stumpp M., Hu M., Casties I., Saborowski R., Bleich M., Melzner F., Dupont S. (2013). Digestion in sea urchin larvae impaired under ocean acidification. Nature Climate Change, 3, 1044–1049.

Talmage S.C., and Gobler C.J. (2010). Effects of past, present, and future ocean carbon dioxide concentrations on the growth and survival of larval shellfish, Proc. Natl. Acad. Sci. U. S. A., 07(40), 17246–17251.

Timmins-Schiffman E., O’Donnell M.J., Friedman C.S., Roberts S.B. (2013). Elevated pCO2 causes developmental delay in early larval Pacific oysters, *Crassostrea gigas*. Marine Biology, 160(8), 1973–1982.

Timmins-Schiffman E., Coffey W.D., Hua W., Nunn B.L., Dickinson G.H., Roberts S.B. (2014). Shotgun proteomics reveals physiological response to ocean acidification in *Crassostrea gigas*. BMC Genomics. doi: 10.1186/1471-2164-15-951.

Timmins-Schiffman E.B., Crandall G.A., Vadopalas B., Riffle M.E., Nunn B.L., Roberts S.B. (2017). Integrating Discovery-driven Proteomics and Selected Reaction Monitoring to Develop a Noninvasive Assay for Geoduck Reproductive Maturation. Journal of Proteome Research, 16(9), 3298–3309.

Vadopalas B., Pietsch T.W., Friedman C.S. (2010) The Proper Name for the Geoduck: Resurrection of *Panopea generosa* Gould, 1850, from the synonymy of *Panopea abrupta* (Conrad, 1849) (Bivalvia: Myoida: Hiatellidae). Malacologia 52(1) 169–173.

Waldbusser G.G., Voigt E.P., Bergschneider H., Green M.A., Newell R.I.E. (2011). Biocalcification in the eastern oyster (*Crassostrea virginica*) in relation to long-term trends in Chesapeake Bay pH. Estuar Coast. 34: 221–231.

Waldbusser G.G., Brunner E.L., Haley B.A., Hales B., Langdon C.J., Prahl F.G. (2013). A developmental and energetic basis linking larval oyster shell formation to acidification sensitivity. Geophysical Research Letters, 40(10), 2171–2176.

Waldbusser G.G., and Salisbury J.E. (2014). Ocean Acidification in the coastal zone from an organism’s perspective: multiple system parameters, frequency domains, and habitats. Annual Rev Marine Sci. 6: 221–247.

Weiss I. M., Tuross N., Addadi L., Weiner S. (2002). Mollusc larval shell formation: Amorphous calcium carbonate is a precursor phase for aragonite, J. Exp. Zool., 293(5), 478–491.

Miller A.W., Reynolds A.C., Sobrino C., Riedel G.F. (2009). Shellfish face uncertain future in high CO2 world: Influence of acidification on oyster larvae calcification and growth in estuaries. PLoS ONE 4(5): e5661. doi:10.1371/journal.pone.0005661

Seung Woo L., Seong Moo H., Cheong Song C. (2006). Characteristics of calcification processes in embryos and larvae of the Pacific oyster, *Crassostrea gigas*. Bulletin of Marine Science, 78(2), 309–317.

