## Supplemental Information 4 for "Dynamic response in the larval geoduck clam proteome to elevated *p*CO_2_"

#
# ABACUS parameter file
# Generated on: 2016Jun30_1144
#

# Name to give the database
dbName=ABACUSDB

# Name of protXML file corresponding to merged/combined results
combinedFile=/net/gs/vol4/shared/nunnlab/search/emmats/TPP/2017_July_24_geolarv/cdhit0.9/lowpH/interact-COMBINED.prot.xml

# The directory that contains the pepXML and protXML files
srcDir=/net/gs/vol4/shared/nunnlab/search/emmats/TPP/2017_July_24_geolarv/cdhit0.9/lowpH

# The name of the file where results will be saved to
outputFile=/net/gs/vol4/shared/nunnlab/search/emmats/TPP/2017_July_24_geolarv/cdhit0.9/lowpH/ABACUS_cdhit0.9_lowpH_output.tsv

# The path the the FASTA formatted file used for the original protein search
# Relative paths are allowed
fasta=/net/gs/vol4/shared/nunnlab/search/emmats/databases/Pgen_larvae_cd0.9.contam

# The minimum PeptideProphet score the best peptide match of a protein must have
maxIniProbTH=0.99

# The minimum PeptideProphet score a peptide must have in order to be even considered by Abacus
iniProbTH=0.50

# E.P.I: Experimental Peptide-probability Inclusion threshold
# If a protein does not contain at least one peptide exceeding this PeptideProphet score, none of the
# peptide evidence for this protein will be considered. This is applied on an experiment by experiment case.
epiTH=0

# The minimum ProteinProphet score a protein group must have in the COMBINED file
minCombinedFilePw=0.90

# If true, Abacus will write ALL protein IDs belonging to a group in the COMBINED file
# Protein IDs starting with ':::' are additional identifiers from the same protein group in
# the COMBINED file. The representative protein for the group does not start with ':::'
verboseResults=false

# The keep the HyperSQL database files that are created after the program is done
keepDB=false

# Should the peptide weights be recalculated in the individual experiment XML files.
# Useful for peptides that are highly degenerate within a single protein groups
recalcPepWts=false

# Spectral count data will be reported in NSAF format.
# NSAF = _N_ormalized _S_pectral _A_bundance _F_actor
# For a detailed explanation of this method refer to this pubmed link:
# http://www.ncbi.nlm.nih.gov/pubmed/20166708
# Abacus reports NSAF values multiplied by a scaling factor. This is done to
# control for numeric underflow (ie: really small numbers). The scaling factor
# that is used is called the NSAF_FACTOR and is reported during runtime in
# case you would like to rescale your data.
asNSAF=true

# If you are using decoy proteins in your searches, specify the first few
# characters of the label indicating decoy proteins here
decoyTag=DECOY_

# Output format that will be produced by this parameter file
output=Default
