## Supplementary figures and images for "Dynamic response in the larval geoduck clam proteome to elevated *p*CO_2_"

### Supplemental Information 6

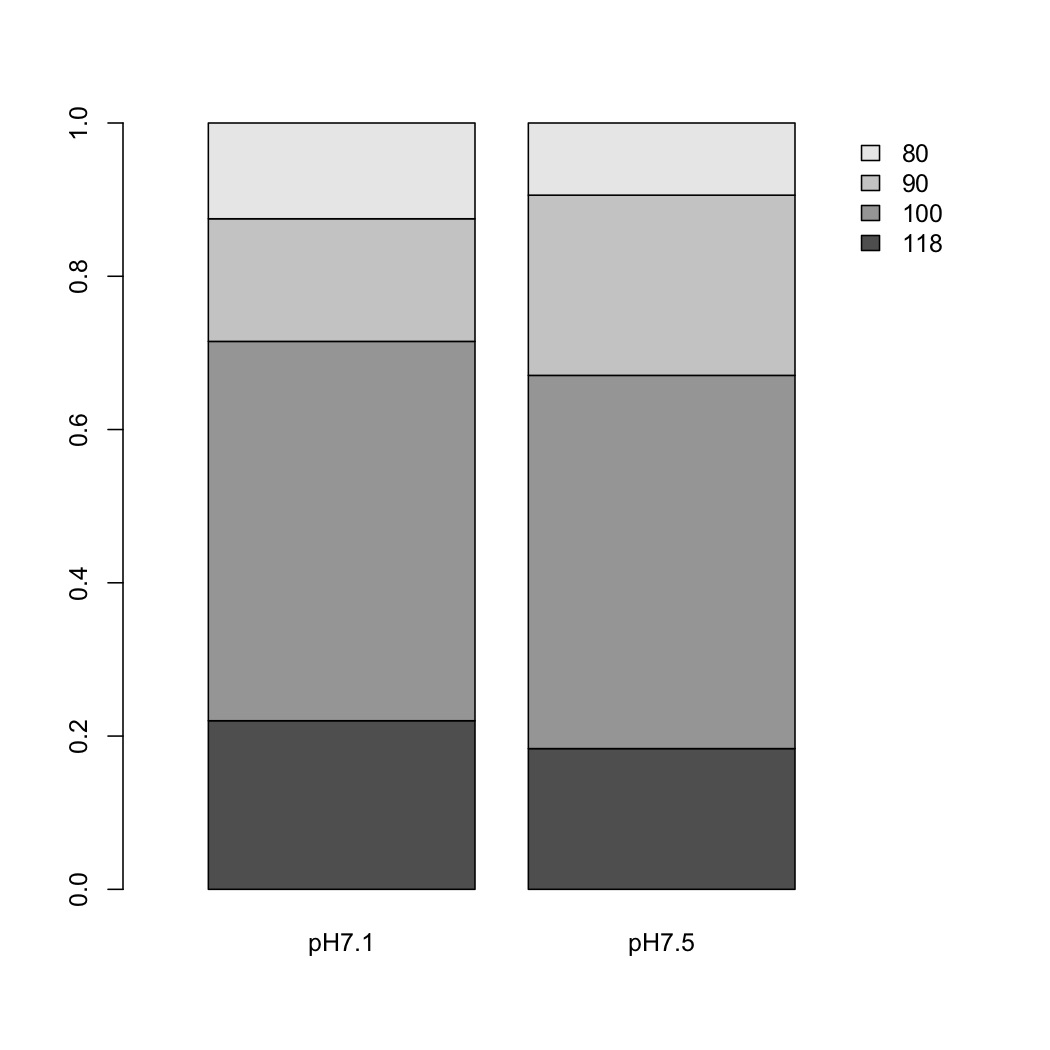
